# Protocols for all-atom reconstruction and high-resolution refinement of protein-peptide complex structures

**DOI:** 10.1101/692160

**Authors:** Aleksandra Badaczewska-Dawid, Alisa Khramushin, Andrzej Kolinski, Ora Schueler-Furman, Sebastian Kmiecik

## Abstract

Structural characterizations of protein-peptide complexes may require further improvements. These may include reconstruction of missing atoms and/or structure optimization leading to higher accuracy models. In this work, we describe a workflow that generates accurate structural models of peptide-protein complexes starting from protein-peptide models in C-alpha representation generated using CABS-dock molecular docking. First, protein-peptide models are reconstructed from their C-alpha traces to all-atom representation using MODELLER. Next, they are refined using RosettaFlexPepDock. The described workflow allows for reliable all-atom reconstruction of CABS-dock models and their further improvement to high-resolution models.

## 1. Introduction

In recent years, peptides have found many successful applications as therapeutic agents and leading molecules in drug design. Consequently, there is a large interest in structural characterization of protein-peptide interactions. Unfortunately, the experimental characterization can be difficult or impossible. Thus, a variety of computational approaches have been developed for structure prediction of the protein–peptide complexes [1]. Computational, or even experimental, structure characterizations often require reconstruction of missing atoms and/or models refinement to a higher resolution. This is also the case for the CABS-dock method [2,3] – a well-established tool for protein-peptide docking.

In this chapter, we present the protocols for all-atom reconstruction of protein-peptide complexes (CABS-dock predictions) from C-alpha trace and their further refinement to higher accuracy models. The software tools used for generating protein-peptide models (CABS-dock), all-atom reconstruction (MODELLER [4]) and refinement (Rosetta FlexPepDock [5]) are shortly presented in Section 2. In section 3, we describe step by step how to use the protocols and discuss their features. Finally, we describe protocol performance on a few example cases in Section 4. The described reconstruction workflow can be applied using input models from CABS-dock, other docking protocols, or sparse experimental data. The advantage of the presented protocols over other protein reconstruction or refinement tools is that they are specifically tailored for protein-peptide complexes. For example, there are many tools that effectively rebuild atomic details from the Cα-trace protein chains (a detailed overview of the available methods can be found in our most recent review [6]). However, most of them cannot be easily used to handle more than one protein chain in the reconstructed structure, and use structural template(s) of the protein backbone to enhance the reconstruction accuracy or to maintain the appropriate interface of protein-peptide interaction. With regard to structure refinement of protein-peptide complexes, the performance robustness and accuracy of the Rosetta FlexPepDock tool has been demonstrated in many studies [7–11].

## 2. Materials

### 2.1. CABS-dock

CABS-dock [2,3] uses an efficient multiscale approach for fast simulation of peptide folding and binding, that merges modeling at coarse-grained resolution with all-atom resolution. The main feature of the CABS-dock method is global flexible docking without a priori knowledge of the binding site and the peptide conformation. However, as an option it is possible to use some knowledge about the interaction interface or peptide conformation in the form of weak distance restraints. CABS-dock is available as a web server [2] and standalone application [3]. CABS-dock has been applied in numerous docking studies, including docking associated with large-scale conformational transitions [12,13]. It has been also extended to modeling protocols that allow for residue-residue contact map analysis of the binding dynamics [14], prediction of protein-protein interaction interfaces [15] and docking using fragmentary information about protein-peptide residue-residue contacts [16].

### 2.2. MODELLER

MODELLER is a program for comparative modeling of protein structures by satisfaction of spatial restraints [4]. The restraints are given as a set of geometrical criteria, which are used to calculate the locations of all non-hydrogen atoms in the protein structure. Smart integration of various MODELLER modules and tailored definition of additional restraints facilitate many other tasks, such as *de novo* modeling of loops, structure refinement, energy optimization or simple all-atom reconstruction from coarse-grained models and addition of hydrogen atoms [17,18].

The required input is an alignment of the query amino acid sequence with known template structures and the corresponding structures data in PDB format. The optional input is a starting structure, e.g. obtained as a result of coarse-grained modeling. The initial structure and templates have to contain at least alpha carbons, but in comparison with all-heavy-atom templates this loses homology information, such as backbone atom distances and side chain dihedrals. The position of a selected type of atoms (e.g. Cα) or chain fragment can be frozen during modeling if necessary.

The MODELLER software is available free of charge to academic use, but a license key valid for a term of 5 years is required. The license key and the software package can be obtained from the website https://salilab.org/modeller/. MODELLER is a command-line only tool and runs on most Unix/Linux systems, Windows, and MacOS. The MODELLER commands are usually provided in a Python script. The user should have basic skills in Python language scripting, but many useful examples of task-dedicated scripts can be found in the *examples* directory.

### 2.3. Rosetta FlexPepDock

FlexPepDock is a high-resolution refinement protocol for modeling of protein-peptide complexes developed within Rosetta framework [19]. FlexPepDock usually requires an approximate pre-defined localization of the binding site. It uses the Monte Carlo-with-minimization approach (MCM) to optimize the rigid body orientation of the peptide, including full flexibility of peptide backbone as well as all receptor side chains. The FlexPepDock is highly effective if the initial peptide conformation is located within 5Å backbone RMSD from the target conformation [5].

The required input is an initial structure of protein-peptide complex. As optional input a reference structure of the protein-peptide complex, such as the native bounded structure, can be provided for RMSD calculations and evaluation of convergence. The initial structure of the complex needs to provide all heavy backbone atoms, including Cβ. The receptor chain coordinates should appear before the peptide coordinates.

The software is available free of charge to academic use, after signing licensing agreement to download the Rosetta package (https://www.rosettacommons.org/software/license-and-download). Rosetta FlexPepDock is also available via a web server at http://flexpepdock.furmanlab.cs.huji.ac.il/.

## 3. Methods

Figure 1 presents the reconstruction and refinement scheme described in this work. The scheme uses ca2all script based on the MODELLER [20] and FlexPepDock [11] protocol.

**Figure 1.**
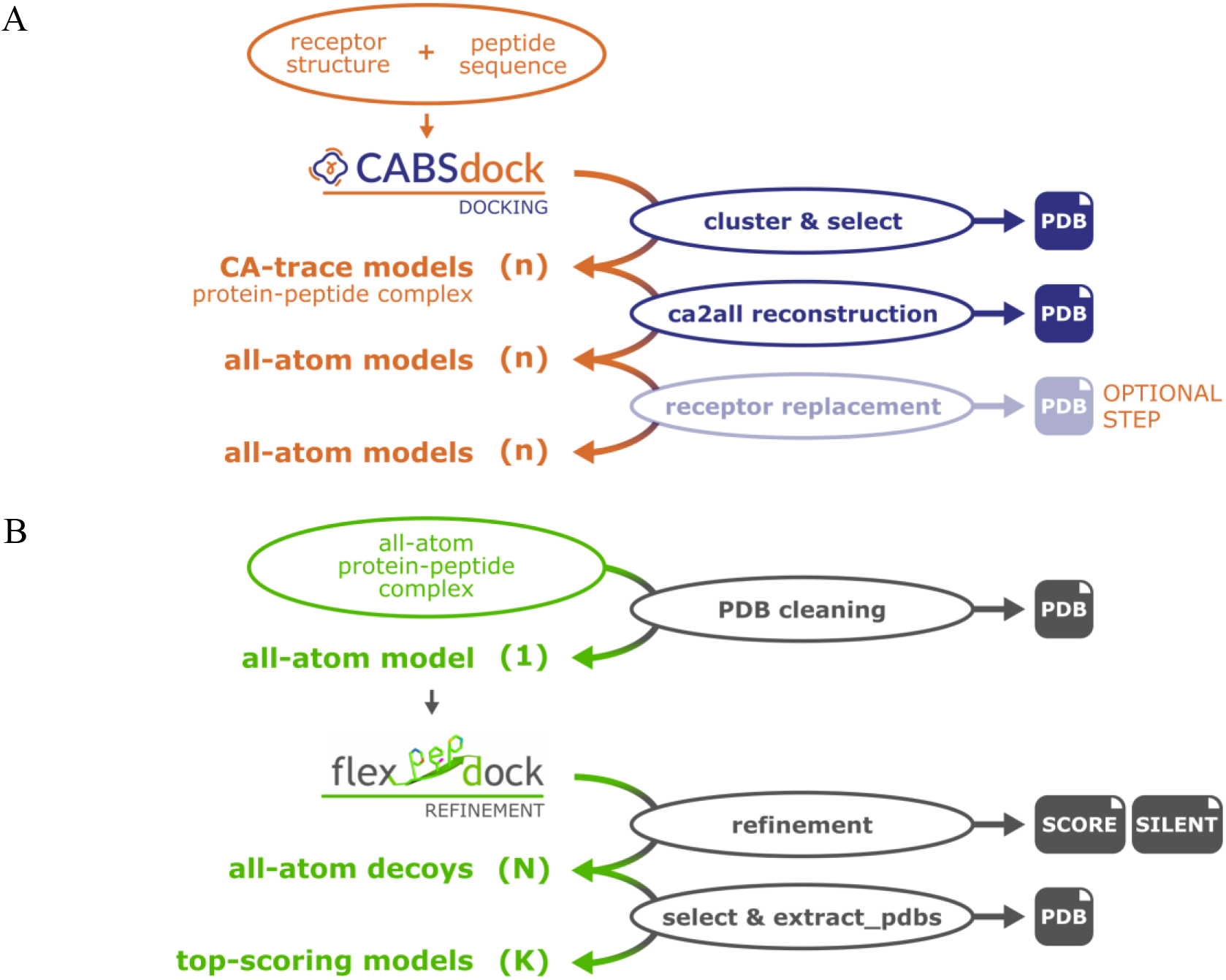
Pipeline of all-atom reconstruction (A) and refinement (B) of protein-peptide complexes. The coarse-grained CABS-dock docking procedure provides n (default n = 10) Cα-traces of top-scoring complex structures. The atomic details for each of these models can be reconstructed using the ca2all.py script based on the MODELLER software. Optionally, the receptor can be replaced by, e.g. the free receptor structure (see the section 4). After removal of clashes in the receptor (using the FlexPepDock prepack protocol), final refinement (using the FlexPepDock refine protocol) is performed. This should include at least 200 independent refinement trajectories (N = 200 optimization runs), of which the top K are extracted and further analyzed (usually K = 10).

### 3.1. Reconstruction of all-atom representations from C-alpha traces using MODELLER

The CABS-dock standalone package [3] offers automatic reconstruction of top-scored models from Cα-trace to atomic resolution using the ***ca2all.py*** script (available from our repository: https://bitbucket.org/lcbio/ca2all/) based on the MODELLER [20] software. Note that CABS-dock releases 0.9.15 and earlier use previous versions of ***ca2all.py*** script, which may lack some of the functionalities described below.

In order to rebuild top-scored models use the --*aa-rebuild* (or -*A*) command-line CABS-dock flag, as in the example below.

| OUTPUT: | Is default? | description |
| --- | --- | --- |
| • model.pdb | YES | output all-atom protein-peptide complex structure |

Note that the ***ca2all.py*** requires MODELLER to be installed. The program offers comprehensive reconstruction and refinement options, as well as some additional user-friendly (fancy) options:

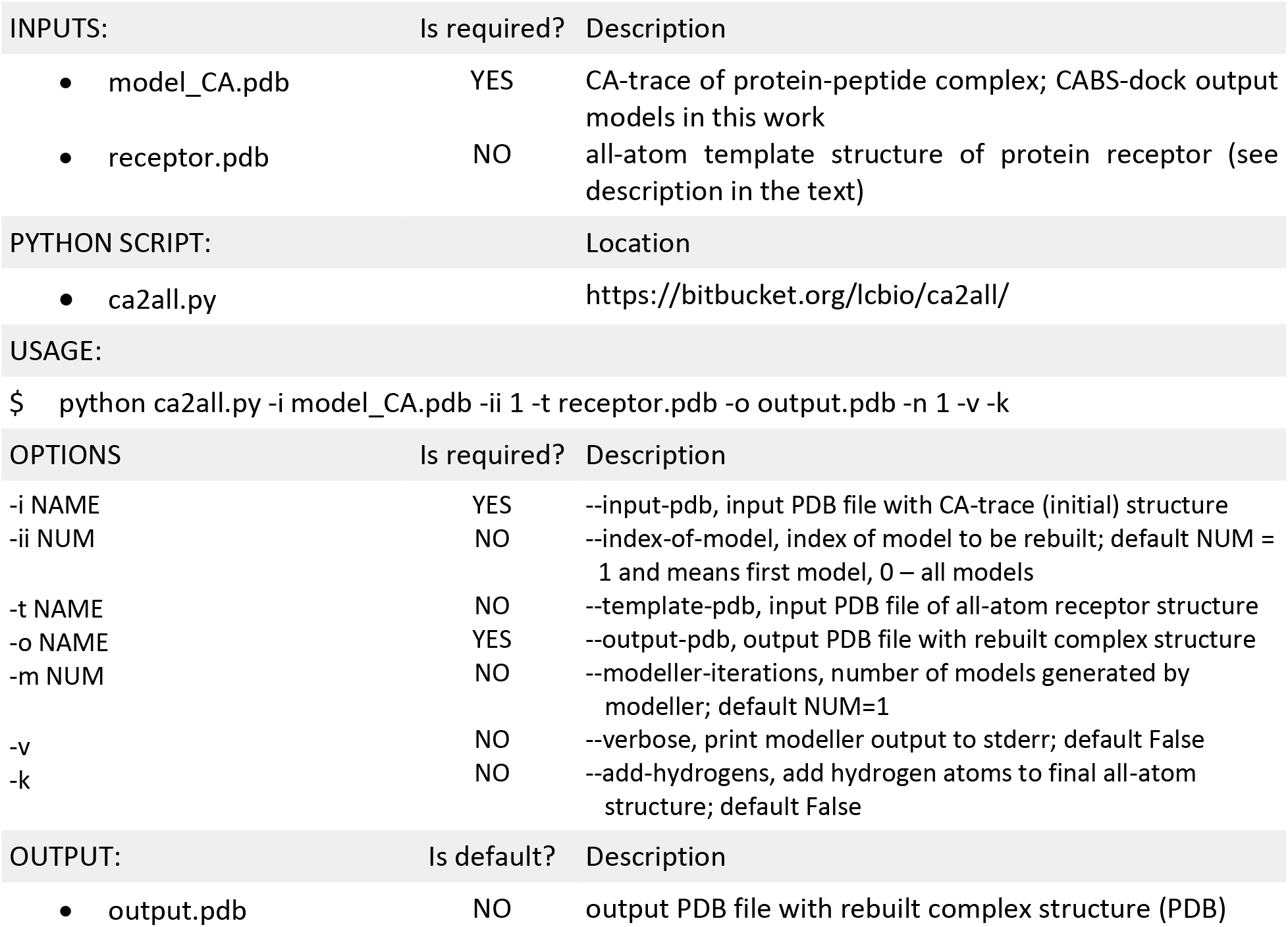

| INPUTS: | Is required? | Description |
| --- | --- | --- |
| • model_CA.pdb | YES | CA-trace of protein-peptide complex; CABS-dock output models in this work |
| • receptor.pdb | NO | all-atom template structure of protein receptor (see description in the text) |

| PYTHON SCRIPT: | Location |
| --- | --- |
| • ca2all.py | https://bitbucket.org/lcbio/ca2all/ |

| OPTIONS | Is required? | Description |
| --- | --- | --- |
| -i NAME | YES | --input-pdb, input PDB file with CA-trace (initial) structure |
| -ii NUM | NO | --index-of-model, index of model to be rebuilt; default NUM = 1 and means first model, 0 – all models |
| -t NAME | NO | --template-pdb, input PDB file of all-atom receptor structure |
| -o NAME | YES | --output-pdb, output PDB file with rebuilt complex structure |
| -m NUM | NO | --modeller-iterations, number of models generated by modeller; default NUM=1 |
| -v | NO | --verbose, print modeller output to stderr; default False |
| -k | NO | --add-hydrogens, add hydrogen atoms to final all-atom structure; default False |

| OUTPUT: | Is default? | Description |
| --- | --- | --- |
| • output.pdb | NO | output PDB file with rebuilt complex structure (PDB) |

The procedure uses an *automodel()* class to build one or more (--*modeller-iterations, -m* option) comparative models based on the CABS-dock-provided final Cα-trace. Note that protein conformations generated by coarse-grained modeling may exhibit some unphysical distortions [21]. This is also the case of the CABS-dock C-alpha trace models obtained from CABS coarse-grained protein model [21]. Those distortions may be not well tolerated by a reconstruction algorithm, since it is not always trivial to keep the desired global topology while maintaining a sound local atomic geometry. We address this issue using appropriate template structures. Based on these, MODELLER creates a set of restraints (mainly distances and angles using heavy atoms and topology). The more detailed the template structure is, the more restrains are provided. For this reason, the reconstruction procedure can use as a template not only the final Cα-chain (from CABS-dock, referred to as *initial*) but also the known all-atom structure of the receptor (referred to as *template*). At the same time, the local geometry of the main chain and the orientation of the side chains for the receptor can be significantly improved by more accurate restraints from the *template* (unpublished results).

The final structure (including alpha carbon atoms) is optimized with the variable target function method with conjugate gradients and refined using molecular dynamics with simulated annealing. Alpha carbon positions are frozen in the first optimization step and freed in the second. By default, the structure returned by ***ca2all.py*** contains all heavy atoms only, but using the --*add-hydrogens* (or -*k*) option will ensure the addition of hydrogen according to geometric criteria (with heavy atoms unchanged). Receptor structures containing chain breaks are not an obstacle to CABS-dock modeling. If a complete sequence of protein is provided, it is possible to rebuild the missing part of the structure using script ***rebuild_loops.py***, which is publicly available in our repository (https://bitbucket.org/lcbio/ca2all/).

Using the ***ca2all.py*** reconstruction protocol it is also possible to reconstruct selected models from the CABS-dock trajectory. By default, this is always the first model in the file, but the --*index-of-model* (or -*ii NUM*) option enables user selection of other models, by specifying *NUM*, the model index (*NUM*=0 means that all models from the trajectory will be processed). If the trajectory is large and you would like to speed up calculations, use our script ***divide_tra*** (available from our repository: https://bitbucket.org/lcbio/ca2all/) that divides the trajectory file into smaller ones and run them parallel on multiple cores.

### 3.2. Refinement of CABS-dock models using Rosetta FlexPepDock

CABS-dock models can be improved by additional structure refinement using FlexPepDock protocol. For the refinement of an individual protein-peptide complex, one can use the online server (available at: http://flexpepdock.furmanlab.cs.huji.ac.il/). Here we will present a variant using the standalone version, *i.e*., a locally installed Rosetta package. Note that Rosetta is written in c++ and needs to be compiled. Detailed description of compilation on different systems can be found at *https://www.rosettacommons.org/docs/latest/build_documentation/Build-Documentation*. After successful installation, some useful scripts for input preparation or results analysis can be found in the directory: ~/Rosetta/tools/protein_tools/scripts/. Some of them needed in this pipeline, such as ***clean_pdb.py*** are described below. The executables of the applications for Rosetta protocols, here ***FlexPepDocking*** or ***extract_pdbs***, can be found in the ~/Rosetta/main/source/bin/ directory, and the database is located in ~/Rosetta/main/databa,e/. In this section we describe how to prepare input files, refine the structure using FlexPepDock and evaluate the results.

#### Prepare an initial complex structure

FlexPepDock protocol requires as input a structure of the protein-peptide complex that includes at least all heavy atoms of the backbone and beta carbon atoms. Coarse-grained CABS-dock models reconstructed to all-atom representation using the MODELLER satisfy this criterion. If CABS-dock models reconstructed with MODELLER contain local structure distortions, replacing the receptor with the experimental structure of the unbound receptor conformation may be helpful (see Discussion in Section 4).

FlexPepDock requires keeping the proper order of coordinates in the file: first the chains of the receptor, then the peptide. It is recommended to keep only the “ATOM” section, although other molecules can be kept if needed (see the Rosetta tutorial “<u>prepare ligand tutorial</u>”). An effective method is to use the Rosetta ***clean_pdb.py*** script, which prints the selected chains in the desired order. The Rosetta refinement protocol only requires the initial structure of the complex, but additional information can also be used if available. The unbound receptor structure is an additional source of rotamers to side chains packing, and the native protein-peptide structure can be used as a reference for the evaluation of refinement results.

| INPUTS: | Is required? | description |
| --- | --- | --- |
| initial_complex.pdb | YES | top-scoring CABS-dock model in all-atom representation |
| ROSETTA SCRIPT: |  | location |
| clean_pdb.py |  | ~/Rosetta/tools/protein_tools/scripts/clean_pdb.py |
| USAGE: |  |  |
| \$ ~/Rosetta/tools/protein_tools/scripts/clean_pdb.py *model.pdb receptor_chain_id peptide_chain_id* | | |
| OUTPUT: | Is default? | description |
| model_AB.pdb | YES | initial complex structure with proper order of chains |

#### Prepack the initial complex structure

Note that the final Rosetta score (*total_score*) takes into account not only the interactions of the protein-peptide interface but also the energy of internal interactions in each monomer. The initial structure of the protein-peptide complex may contain local distortions, especially side chain clashes or backbone distortions in the receptor structure that will result in high Rosetta’s energy penalties and will not reliably identify best refined models. In order to avoid the effects of non-uniform conformational background in non-interface regions, Rosetta repacking with the -*flexpep_prepack* flag is recommended. If several models are generated from a specific starting model, it is suggested to use the same prepacked structure as starting point. In Section 4, we describe a modeling option in which we replace protein receptor coordinates in CABS-dock models with experimentally determined unbound receptor structures (see Note 1 and Section 4). This can be helpful if distortions at the backbone level occur that are not removed by side chain repacking.

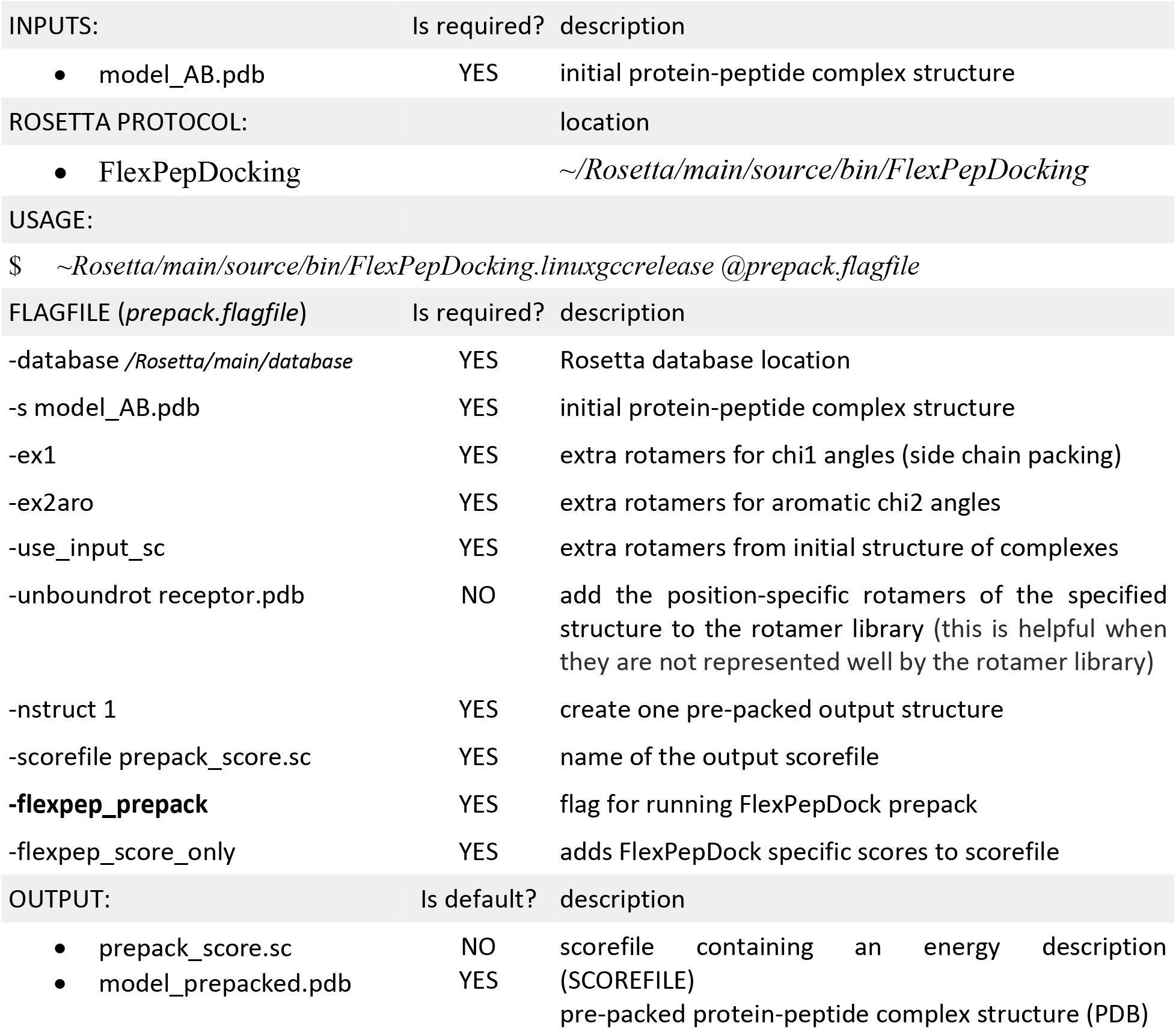

#### Refine the prepacked complex structure

FlexPepDock refinement protocol uses an iterative Monte Carlo with energy minimization scheme to optimize the position of the peptide backbone relative to the receptor and the orientation of the side chains in the interaction interface. It is recommended to generate at least 200 of refined models (-*nstruct* flag), whereas each of them is an independent event. Additional improvement can be achieved through initial pre-optimization at the coarse-grained stage (Rosetta centroid representation) by using -*lowres_preoptimize* flag.

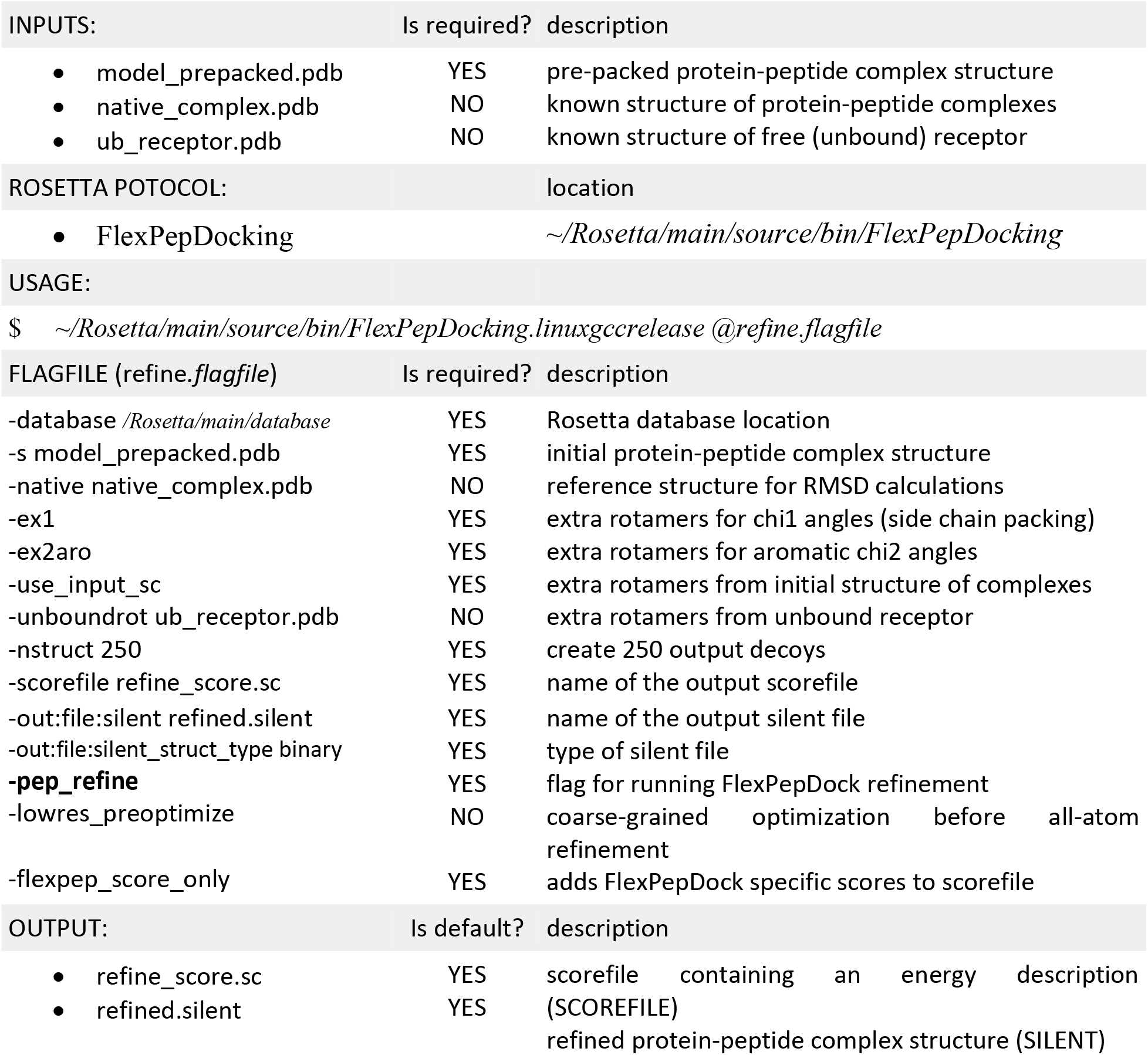

#### Analysis of refinement results

The refined output models are saved to a file (-*out:file:silent* flag). The silent file is a Rosetta output format that is used to store compressed ensembles of structures. Each frame in a silent file has a unique identifier, which is called the decoy-tag. The unique decoy-tag is located at the end of each line that belongs to the respective frame and enables to identify and extract selected frames. The second output file is a score file (-*scorefile* flag) that contains the energy description of each of the output models and various root-mean-square deviation (RMSD) parameters in relation to reference complex structure (if provided) or to initial complex structure. This file allows to analyse the energy landscape sampled during the refinement procedure (by ploting *score* vs. *rmsd*), or select top-best models (sorted according to Rosetta energy criterion). To evaluate FlexPepDock refinement results it is recommended to use interface_score (*I_sc*), reweighted_score (*reweighted_sc*, the weighted sum of total *score*, *I_sc* and *pep_sc*) rather than the total score *rmsBB_if* (peptide backbone interface RMSD). Selected top-scoring models can be extracted into pdb format by using Rosetta ***extract_pdbs*** protocol.

|  |  |  |  |  |  |  |  |  |  |  |  |
| --- | --- | --- | --- | --- | --- | --- | --- | --- | --- | --- | --- |
| SCORE: | score | ... | reweighted_sc | ... | I_sc | ... | pep_sc | ... | rmsBB_if | ... | decription |
| SCORE: | -141.009 | ... | -117.833 | ... | 6.454 | ... | 16.722 | ... | 4.117 | ... | model_refined_0001 |
| SCORE: | -215.024 | ... | -199.562 | ... | 4.598 | ... | 10.864 | ... | 2.006 | ... | model_refined_0155 |

Example of sorting refined decoys by *reweighted_sc* and selecting 10 top-scoring models

|  |  |  |
| --- | --- | --- |
| INPUTS: | Is required? | description |
| • refine_score.sc | YES | score file |
| OUTPUT: | Is default? | description |
| • top10-tags | NO | 10 top-scoring models |

Example of extracting PDB coordinates for selected decoy-tags

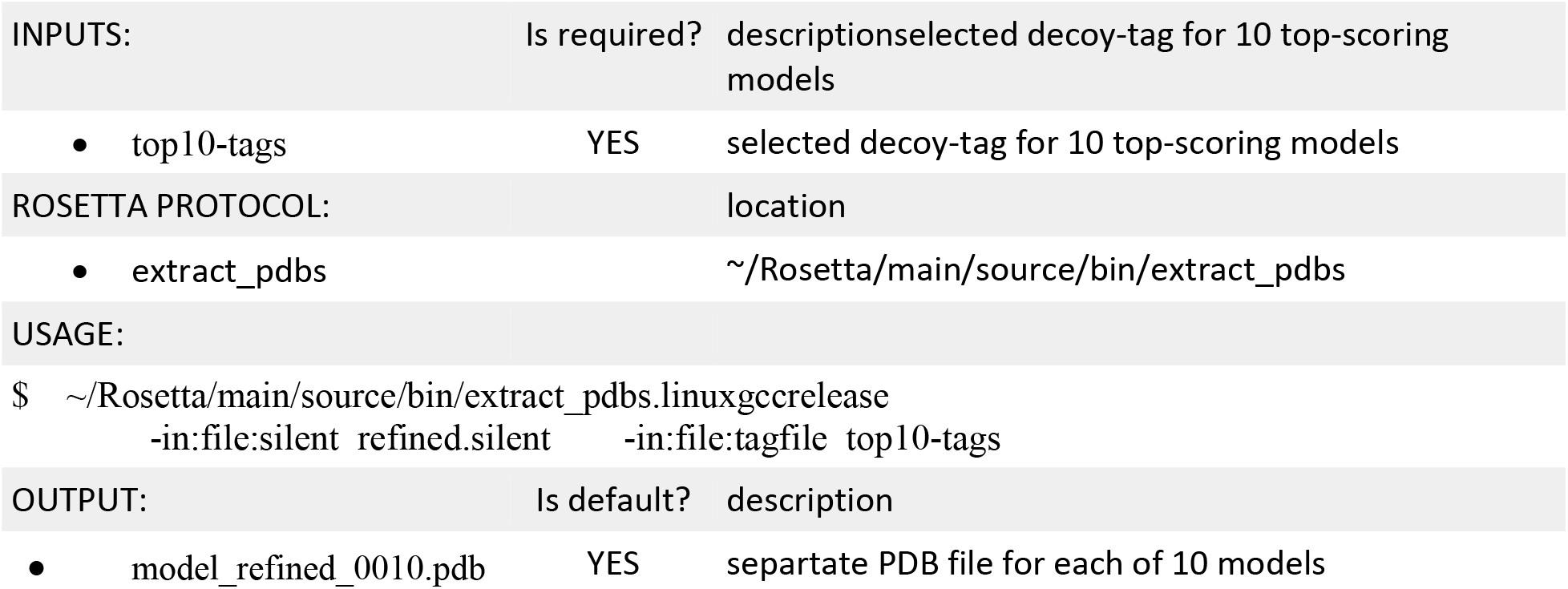

## 4. Case Studies

We detail here the implementation and results of the above described protocol for the atomic reconstruction and refinement. The starting point are CABS-dock predictions of four different protein-peptide complexes from our previous study by Blaszczyk *et al*.[16]. For each protein-peptide complex, we reconstructed 10 top-scored CABS-dock models to all-atom structure using the ***ca2all*** script. Next, in each reconstructed model we replaced the receptor structure with the unbound receptor structure (see Note 1). Note that this is an optional step in the reconstruction and refinement procedure described in this work. Introduction of conformational flexibility in the receptor can be important to identify binding sites in cases where the receptor changes considerably upon binding. However, introduction of conformational flexibility might also lose features of the receptor that are crucial to allow the high-resolution step to identify the native conformation among other false positive minima in the energy landscape. Thus, by replacing the receptor with the solved structure of the unbound receptor conformation, we allow both the detection of cryptic binding sites in the first step, and the refinement starting from an informative crystal structure in the second step. The peptide coordinates were taken directly from CABS-dock modeling. In order to remove all possible internal clashes and to create a uniform energy background, the receptor unbound structure was prepacked using the Rosetta -*flexpep_prepack* protocol. Prepacking was performed once, and the same receptor structure was used for all simulations. Finally, each resulting model was further refined with the *FlexPepDock refinement* protocol, and the 10 top-scored models were selected for quality assessment. Figure 2 shows comparison of the top-scoring refined model with CABS-dock models and experimental reference structures. As presented in the Figure, FlexPepDock refinement can significantly improve the modeling accuracy to sub-Ångstrom level, as measured by i-RMSD (interface root mean square deviation from the reference experimental structure). Motivated by these promising results we are now applying this protocol on large and representative benchmark set.

**Figure 2.**
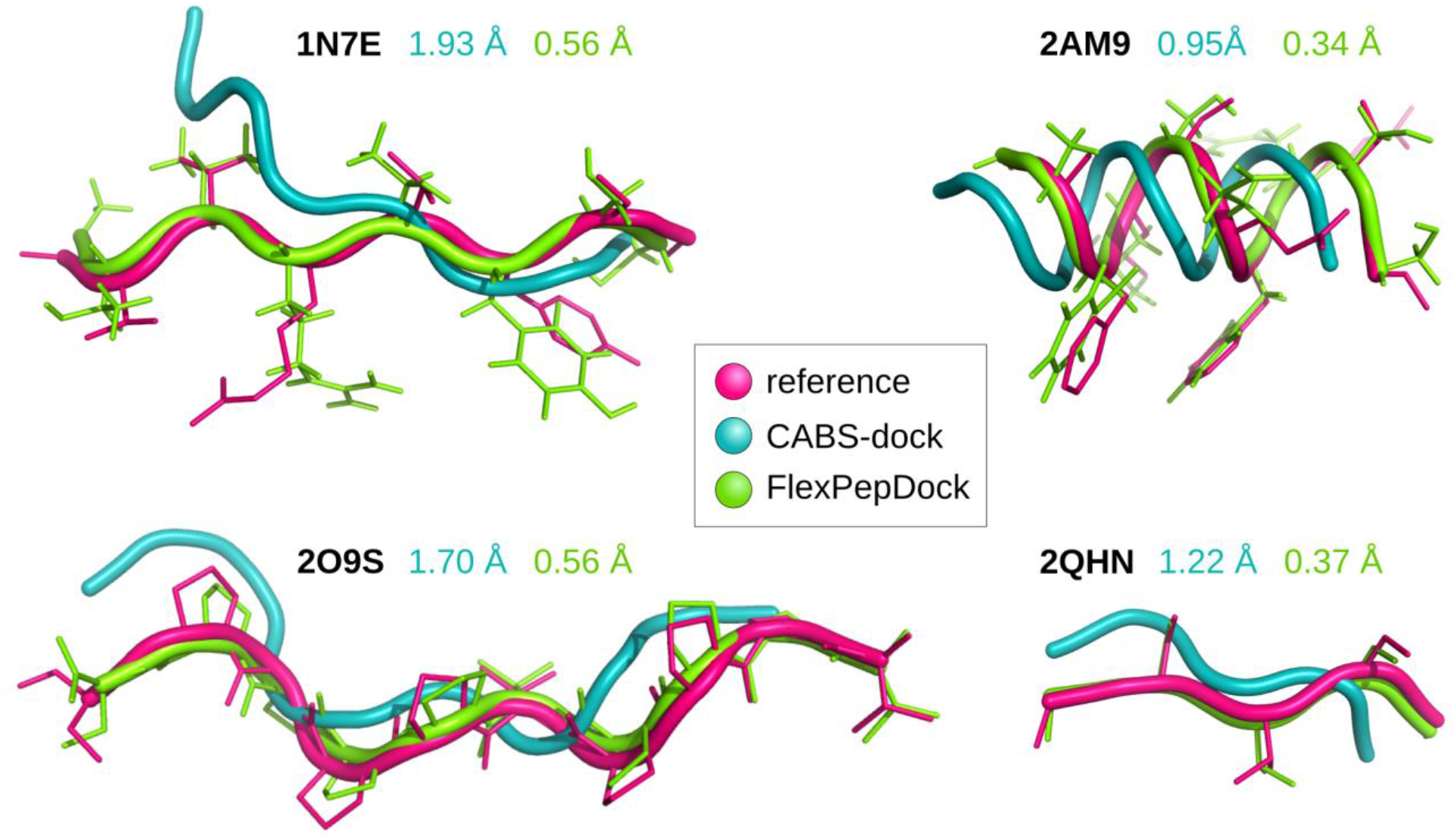
Comparison of CABS-dock models before and after FlexPepDock refinement, to experimental structures. The figure presents four example cases of protein-peptide complexes (1N7E, 2O9S, 2AM9 and 2QHN). For each complex, only peptide conformations are shown (after superimposition of the receptor structures). Reference experimental structures are shown in pink, CABS-dock models before refinement in blue and after in green. The numbers in the corresponding colors indicate i-RMSD values.

## 5. Notes

### Note 1

In order to replace the structure of receptor, you should first align the CABS-dock model of the complex and the unbound receptor using, e.g., Theseus[22] (or PyMOL):

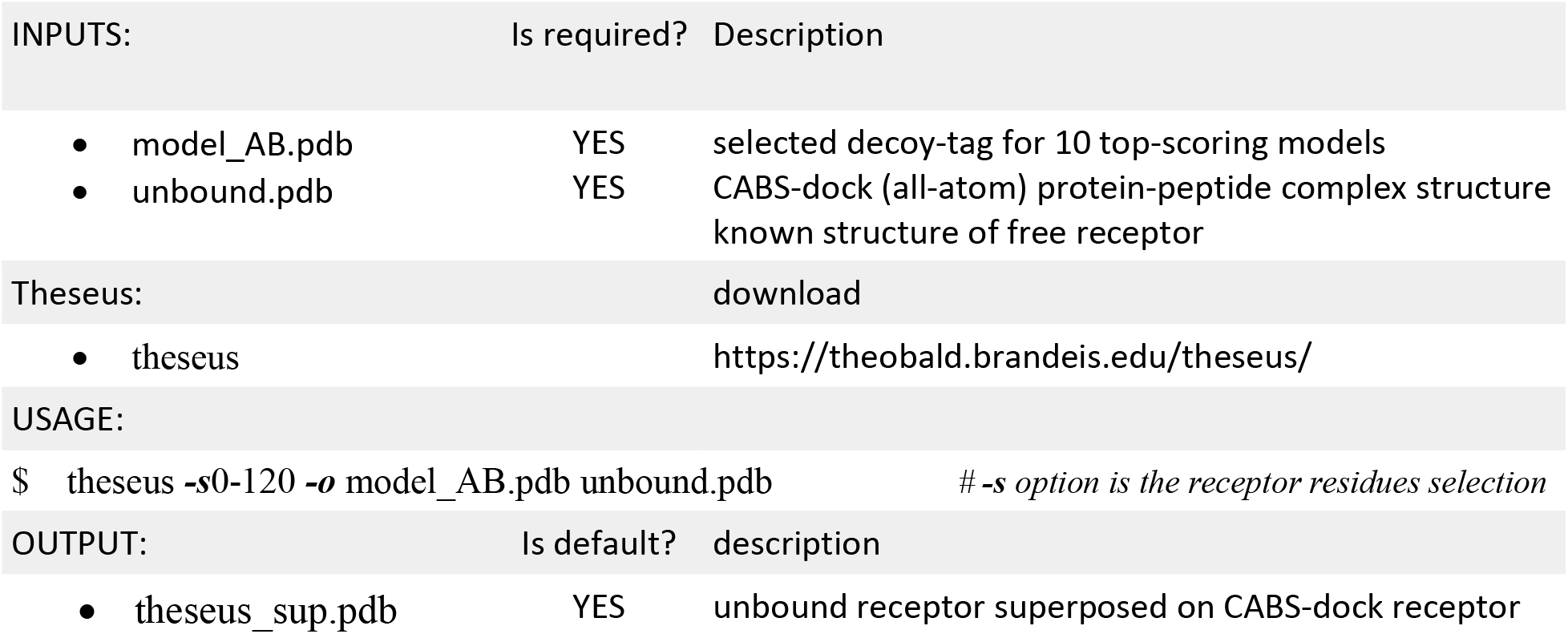

In the next step you need to cut the peptide coordinates from the *model_AB.pdb* file, which is possible using the already known Rosetta ***clean_pdb.py*** script and then add peptide to the free receptor *theseus_sup.pdb* file.

|  |  |  |
| --- | --- | --- |
| USAGE: |  |  |
| \$ ~/Rosetta/tools/protein_tools/scripts/clean_pdb.py model_AB.pdb B | # cut peptide (chain B) | |
| \$ cat theseus_sup.pdb model_AB_B.pdb > model.pdb | # paste unbound receptor and peptide | |
| OUTPUT: | Is default? | description |
| • model.pdb | YES | free receptor and CABS-dock modeled peptide |

## Acknowledgments

A.E.B-D, A.Ko. and S.K. received funding from NCN Poland, Grant MAESTRO2014/14/A/ST6/00088. O.S-F. and A.Kh. received funding from the ISF, Grant 717/17.

